# Cortical activity evoked by voice pitch changes: a combined fNIRS and EEG study

**DOI:** 10.1101/2020.08.24.264275

**Authors:** Kurt Steinmetzger, Esther Megbel, Zhengzheng Shen, Martin Andermann, André Rupp

## Abstract

Numerous studies have investigated the cortical representation and time course of responses to auditory signals with and without pitch. However, little is known regarding the responses evoked by pitch changes. To study the spatial as well as temporal characteristics of cortical activity elicited by voice pitch changes, combined functional near-infrared spectroscopy (fNIRS) and electroencephalography (EEG) data were obtained from normal-hearing listeners presented with continuous vowel sequences in which the prosodic contours either varied between the individual vowels or were the same throughout. The fNIRS topographies and the EEG source reconstructions indicated additional right-hemispheric activity anterior to the primary auditory cortex and in the superior temporal sulcus for sequences with variable prosody. Additionally, the time courses of the fNIRS signals showed that activity was more sustained in response to sequences with changing prosodic contours. The EEG data revealed a similar pattern for the P2 amplitude, which was smaller during the second half for blocks with fixed prosody, whereas the P1 was consistently larger for sequences with variable prosody. Moreover, the sources of the P2, but not the P1, were lateralised to the right, suggesting that the hemispheric asymmetry in the processing of voice pitch changes develops over time.

## I. INTRODUCTION

Pitch changes in auditory signals transmit crucial information, such as prosodic variations in speech and melody in music. Pitch information also promotes the formation of separate auditory streams (Oxenham, 2008), which is an important reason for why it helps understanding speech in noise (Cullington and Zeng, 2008; Steinmetzger and Rosen, 2015). The restricted access to pitch information, in turn, is a critical limitation when listening through a cochlear implant (Macherey and Carlyon, 2014; Steinmetzger and Rosen, 2018).

Previous research has identified a region close to the anterolateral border of the primary auditory cortex, referred to as ‘pitch centre’, that is activated by auditory input giving rise to a pitch percept (Bendor and Wang, 2005; Patterson et al., 2002; Penagos et al., 2004). The time course of pitch-related neural activity has been investigated by many electrophysiological studies that have consistently found the transient N1 component (e.g. Andermann et al., 2017; Chait et al., 2006; Krumbholz et al., 2003; Ritter et al., 2005; Schönwiesner and Zatorre, 2008) as well as the sustained field in magnetoencephalography (MEG) data (Gutschalk et al., 2002; Gutschalk et al., 2004) to increase in response to sounds with a pitch. Using continuous stimulus designs, where the pitch onset occurred after the initial sound onset and without affecting the acoustical properties of the ongoing stimulus (Chait et al., 2006; Gutschalk et al., 2004; Krumbholz et al., 2003; Schönwiesner and Zatorre, 2008) and dipole modelling techniques of MEG data (Andermann et al., 2017; Gutschalk et al., 2002; Ritter et al., 2005), these studies were able to distinguish between pitch responses and sound energy onset responses that are generated slightly further posterior, in planum temporale.

Regarding the processing of pitch changes, evidence from lesion and neuroimaging studies (Boemio et al., 2005; Johnsrude et al., 2000; Patterson et al., 2002) suggests that non-primary regions in the right hemisphere are critically involved. In particular, Patterson et al. (2002) have shown that while tones with a fixed pitch mainly evoke activity in primary auditory cortex, pitch changes elicit additional activity in the superior temporal sulcus (STS) as well as in planum polare, a region slightly anterior to primary auditory cortex. Yet, only very little electrophysiological results concerning the processing of pitch changes are available. For example, the MEG data of Rupp and Uppenkamp (2005) showed that both the N1m and the P2m amplitude in response to pitch changes were enhanced compared to tone sequences with a fixed pitch. Patterson et al. (2016) reported similar MEG results, with increased N1m and especially P2m amplitudes following pitch changes, the latter of which originated from the anterior part of the primary auditory cortex. Moreover, both of these studies as well as the results of Patterson et al. (2002) are based on discrete, music-like pitch changes, rather than continuous pitch changes typical for speech. Here, the aim was to continue this line of research by investigating cortical responses to voice pitch changes as they appear in natural speech.

To investigate spatial as well as temporal aspects of the responses evoked by voice pitch changes, we obtained combined functional near-infrared spectroscopy (fNIRS) and electroencephalography (EEG) data from normal-hearing listeners. Additionally, this approach allowed for the mutual validation of the results, particularly regarding the fNIRS topographies and EEG source reconstructions. Briefly summarised, fNIRS measurements are based on the relatively low absorption of infrared light by biological tissue, resulting in a so-called optical window into the brain (Pinti et al., 2018; Scholkmann et al., 2014). Source optodes placed directly on the scalp emit infrared light towards the brain, while detector optodes positioned at scalp sites nearby record the amount of light that has passed the cortical area in between the two sensors. The more active a given cortical region is, the more light will be absorbed, since brain activity causes an inflow of oxygenated blood containing higher concentrations of red oxyhaemoglobin (HbO). This increase in HbO concentration over time, and the concurrent smaller decrease of the concentration of de-oxygenated haemoglobin (HbR), can be measured using fNIRS.

The stimuli used in this experiment were German vowels concatenated into continuous sequences. As in a previous fNIRS-EEG study (Steinmetzger et al., 2020), long stimulus blocks were used to maximise the fNIRS responses, making adaptation processes likely. The pitch contours within each sequence were either the same for each individual vowel (fixed prosody) or varied between vowels (variable prosody). We hypothesised that both conditions would evoke activity in auditory cortex as well as associated regions in the superior temporal lobe, since vowels have been shown to elicit prominent bilateral activity in STS (Uppenkamp et al., 2006), but that sequences in which the pitch contours changed would evoke additional right-lateralised activity beyond primary auditory cortex. It was furthermore assumed that this additional activity in the right hemisphere would be reflected in larger P2 amplitudes in the EEG waveforms.

## II. METHODS

### A. Participants

Twenty subjects (10 females, 10 males; mean age 26.2 years, standard deviation 8 years) participated in this experiment. All subjects were German native speakers and had audiometric thresholds of less than 20 dB hearing level at octave frequencies between 125 and 8000 Hz. All participants reported that they are right-handed and have no history of neurological or psychiatric illnesses. Written consent was obtained prior to the experiment and the study was approved by the local research ethics committee (Medical Faculty, University of Heidelberg).

### B. Stimuli

The stimulus materials used in this study were recordings of the German vowels /a/, /e/, /i/, /o/ and /u/ spoken by an adult male German talker. The recordings were made in an anechoic room and digitised with 24-bit resolution and a 48-kHz sampling rate, using a condenser microphone (Brüel & Kjær, type 4193, Nærum, Denmark) and an RME Babyface audio interface (Haimhausen, Germany). Each vowel was cut at zero-crossings right before vowel onset, limited to a length of 800 ms using a 50-ms Hann-windowed offset ramp, and high-pass filtered at 50 Hz (zero-phase-shift 3^rd^-order Butterworth).

Subsequently, the original *F*0 contours of the vowels were manipulated with the STRAIGHT vocoder software (Kawahara and Irino, 2005) implemented in MATLAB (MathWorks, Natick, MA, USA), which allows to alter the prosodic properties of the stimulus materials without affecting their spectral envelope. Each of the five vowels was re-synthesised with two different mean fundamental frequencies (*F*0s; 80 Hz and 120 Hz) and five different prosodic contours (flat, rising straight, falling straight, rising curved, and falling curved), resulting in a final set of 50 stimuli. For the non-flat contours, the *F*0 increased or decreased by a perfect fifth relative to the mean *F*0. The mean *F*0 was always specified to be the mid-point of the non-flat contours, such that the maximum and minimum *F*0 values were equally far above and below the mean. Finally, the stimuli were low-pass filtered (zero-phase-shift 1^st^-order Butterworth) at 3.5 kHz – to match the frequency response of the Etymotic Research ER3 headphones (Elk Grove Village, IL, USA) used in a corresponding MEG experiment – and normalised to a common root-mean-square level.

Examples of the stimuli are shown in Fig. 1A. The narrow-band spectrograms depicted in the upper half of the figure demonstrate that the spectrum of the example vowel (/e/, mean *F*0 = 120 Hz) is indeed very similar across the five prosodic versions, while the waveforms differ markedly. To visualise the different prosodic contours, the lower half of the plot shows spectrographic representations of summary autocorrelation functions (SACFs; Meddis and Hewitt, 1991; Meddis and O’Mard, 1997). The individual autocorrelation functions were calculated for the low-pass filtered (2^nd^-order Butterworth, cut-off 1 kHz) outputs of 22 gammatone filters with centre frequencies ranging from 0.2–4 kHz and summed together into SACFs. This procedure was applied with a step size of 1 ms and a Hann-window size of 5 ms to yield spectrographic representations of the SACFs across the duration of the stimuli. The time lag of the first peak in the SACF spectrograms represents the *F*0 of the stimuli.

**Figure 1.**
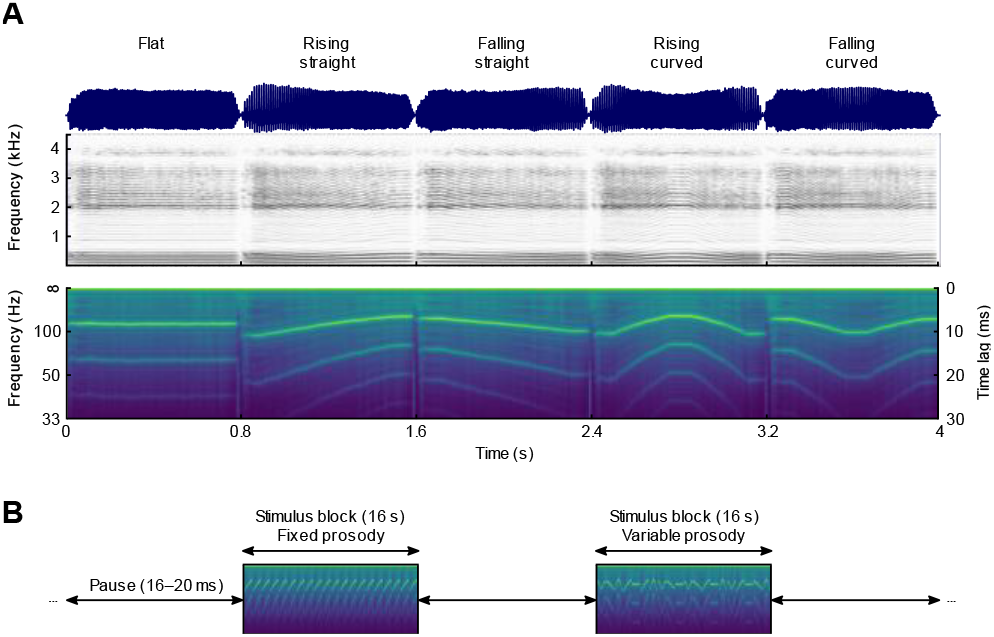
Example stimuli and experimental design. A) Waveforms, narrow-band spectrograms, and summary autocorrelation function spectrograms of the vowel /e/ processed to have five different prosodic contours. B) The individual vowels were concatenated into blocks of 20 vowels lasting 16 s and alternated with pauses lasting 16–20 s.

### C. Experimental design and procedure

As haemoglobin concentration changes evolve over the course of several seconds, a block design with long stimulation periods and equally long pauses was used to maximise the size of the experimental effects. The individual vowels were thus concatenated into blocks of 20 stimuli with a total duration of 16 s. These stimulus blocks were followed by pauses with random durations ranging from 16–20 s. The experiment consisted of two main conditions (fixed prosody and variable prosody), each formed of five subconditions. In the fixed prosody condition, the vowels within each block all had prosodic contours that were either flat, rising straight, falling straight, rising curved, or falling curved. In the variable prosody condition, where the contours varied between the vowels within each block, the five subconditions were rising, falling, straight, or curved contours, as well as a mixture of all five contour types. Each participant was presented with 10 blocks in each subcondition, i.e. 50 blocks in each main condition. In each block, the same vowel with the same mean *F*0 was presented throughout and there was one block for each of the ten combinations of vowel type and mean *F*0 in each subcondition, to ensure that all combinations occurred equally often across subconditions. The order of the blocks as well as the order of the prosodic contours within each block of the variable prosody condition were both randomised without any constraints. The experiment thus consisted of 100 stimulus blocks, framed by 101 pauses, amounting to a total duration of about 57 mins. As each block contained 20 individual vowels, this design resulted in 1000 EEG trials (50 blocks x 20 vowels) in both the fixed and variable prosody conditions.

The experiment took place in a sound-attenuating and electrically shielded room, with the participant sitting in a comfortable reclining chair during data acquisition. Throughout the experiment, the light was dimmed to the lowest level to minimise ambient light from interfering with the fNIRS recordings. There was no behavioural task, but pauses were inserted about every 15 mins to ensure the vigilance of the subjects. The stimuli were with 24-bit resolution at a sampling rate of 48 kHz using an RME ADI-8 DS sound card and presented via Etymotic Research ER2 earphones attached to a Tucker-Davis Technologies HB7 headphone buffer (Alachua, FL, USA). The presentation level was set to 70 dB SPL, using an artificial ear (Brüel & Kjær, type 4157) connected to a corresponding measurement amplifier (Brüel & Kjær, type 2610).

### D. fNIRS recording and analysis

fNIRS signals were recorded with a continuous-wave NIRScout 16×16 system (NIRx Medizintechnik, Berlin, Germany) at a sampling rate of 7.8125 Hz. Eight source optodes and eight detector optodes were placed symmetrically over each hemisphere by mounting them on an EEG cap (EasyCap, Herrsching, Germany). The source optodes emitted infrared light with wavelengths of 760 and 850 nm. To avoid interference between adjacent sources, only a single source optode per hemisphere was illuminated at a given time. The chosen optode layout was devised to optimally cover the auditory cortex and associated areas, resulting in 22 measurement channels per hemisphere, of which 20 had a standard source-to-detector distance of about 30 mm, while the remaining 2 had a shorter spacing of about 15 mm. The optode and reference positions for each individual subject were digitised with a Polhemus 3SPACE ISOTRAK II system (Colchester, VT, USA) before the experiment.

The data were pre-processed using the HOMER2 toolbox (version 2.8; Huppert et al., 2009) and custom MATLAB code. The raw light intensity signals were first converted to optical density values and then corrected for motion artefacts. A kurtosis-based wavelet algorithm with a threshold value of 3.3 (Chiarelli et al., 2015) was used to identify and correct motion artefacts by rejecting spectral components of the signal rather than time segments. Measurement channels with poor signal quality were then excluded from further analysis based on their scalp coupling index (SCI; Pollonini et al., 2014). SCIs were computed by filtering the optical density signals of both wavelengths between 0.5–2.5 Hz (3^rd^-order low-pass and 5^th^-order high-pass Butterworth filters, applied forwards and backwards), to emphasise the heart-beat related signal fluctuations, and correlating the filtered signals. Channels with correlation coefficients below 0.5 were excluded (mean = 1.7 channels/subject, max. 10 per subject), as this indicates a poor contact between optodes and scalp. As we did not pre-select the subjects, the sample included several participants with dark, long hair. Thus, a lower SCI threshold than the one suggested by Pollonini and colleagues (0.75) was used to limit data loss. Next, the motion-corrected signals of the remaining channels were band-pass filtered between 0.01–0.5 Hz (same filter types as for the SCI above), to isolate the task-related neural activity, and subsequently converted to concentration values based on the modified Beer-Lambert law (Scholkmann et al., 2014). The differential path length factors required for the conversion were determined based on the wavelengths of the light and the age of the subject (Scholkmann and Wolf, 2013).

Secondly, the pre-processed data were statistically evaluated and topographically visualised with SPM-fNIRS (version r3; Tak et al., 2016). Based on the principles of the general linear model (GLM), the SPM framework tests how closely the stimulus-evoked signal changes resemble a canonical haemodynamic response function (HRF). In SPM-fNIRS, the optode positions of each subject were first transformed from subject space to MNI space, after which they were probabilistically rendered onto the ICBM-152 cortical template surface. This was achieved by applying a least-squares approach in which a set of digitised reference points distributed across the whole head (4 external fiducials and 17 positions of the extended international 10-20 system) was aligned to template values (Singh et al., 2005). The signals were then temporally smoothed using the shape of the canonical HRF waveform (‘pre-colouring’) to avoid autocorrelation issues when estimating the model (Worsley and Friston, 1995). The data of the individual subjects were statistically modelled by convolving the continuous signals obtained from each long channel with separate regressor functions representing the fixed and variable prosody conditions. The standard SPM double-gamma function was used as canonical HRF and convolved with 16-s boxcar functions following the onsets of the stimulus blocks. The HbO data were modelled with positive HRFs, while the concentration changes were assumed to be negative for the HbR analysis. To allow the time course of the measured concentration changes to vary slightly, the temporal and spatial derivatives of the canonical HRF were included as additional regressors (Plichta et al., 2007). Furthermore, the first component of a principal component analysis of the pre-processed signals of the four short channels was used as an additional nuisance regressor, as this serves to estimate and remove the so-called global scalp-haemodynamic component (Sato et al., 2016), i.e. the superficial signal component. After estimating the HbO and HbR GLMs for each subject, contrast vectors were defined in which the regressors of the five subconditions in the fixed or variable prosody conditions were set to 1, whereas the regressors representing the other five subconditions, as well as all the derivatives and the global scalp component were set to 0 to statistically control for their effects. Group-level statistics were then computed for each long channel by testing whether the subject-level β weights in the fixed or variable prosody conditions were significantly greater than 0. Non-parametric right-tailed Wilcoxon signed-rank tests were used, as Kolmogorov-Smirnov tests indicated that the β weights for the HbO and HbR data of many long channels had non-normal distributions. When comparing the two conditions, the regressors of the variable prosody condition were set to 1, while the regressors of the fixed prosody condition were set to −1, as we expected stronger activity in the former condition. Non-parametric right-tailed Wilcoxon rank-sum tests were used for this comparison and the regressors representing the derivatives and global scalp component were set to 0, as before.

A customised version of the SPM-fNIRS plotting routine was devised to topographically visualise the optode and channel positions as well as the functional activations. The MNI-transformed optode and channel locations as well as the measured channel lengths are shown in Fig. 2. The short channels were omitted in the plots of the functional activations (Fig. 3A & 4A) as they are assumed to not reflect any cortical responses. In the plots depicting the HRFs (Fig. 3B & 4B), the waveforms were averaged from −2–32 s around block onset and baseline corrected by subtracting the mean amplitude in the pre-stimulus window from each sample point. The corresponding HRFs are shown both after the pre-processing (‘Total HRF’) and again after regressing out the contribution of the short channels and pre-colouring the signals (‘Cortical HRF’), to illustrate the effect of removing the non-cortical signal component. Additionally, the β-weighted canonical HRFs (‘Model HRF’) are shown to demonstrate how well the cortical HRFs can be explained by the HRF models.

**Figure 2.**
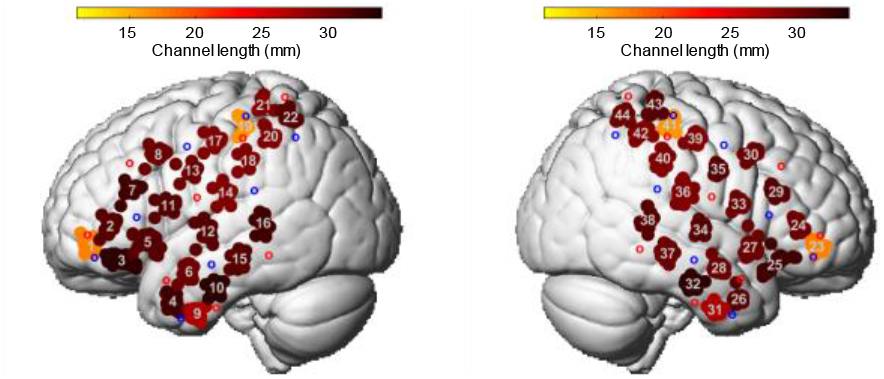
fNIRS measurement layout. The group-averaged MNI positions of the source and detector optodes are shown as red and blue circles, respectively. The average positions of the resulting measurement channels are indicated by the grey numbers. The point cloud around each channel signifies the positions of the individual subjects. The colour of the cloud shows the group-averaged channel length, i.e. the mean distance between the respective source-detector pair.

**Figure 3.**
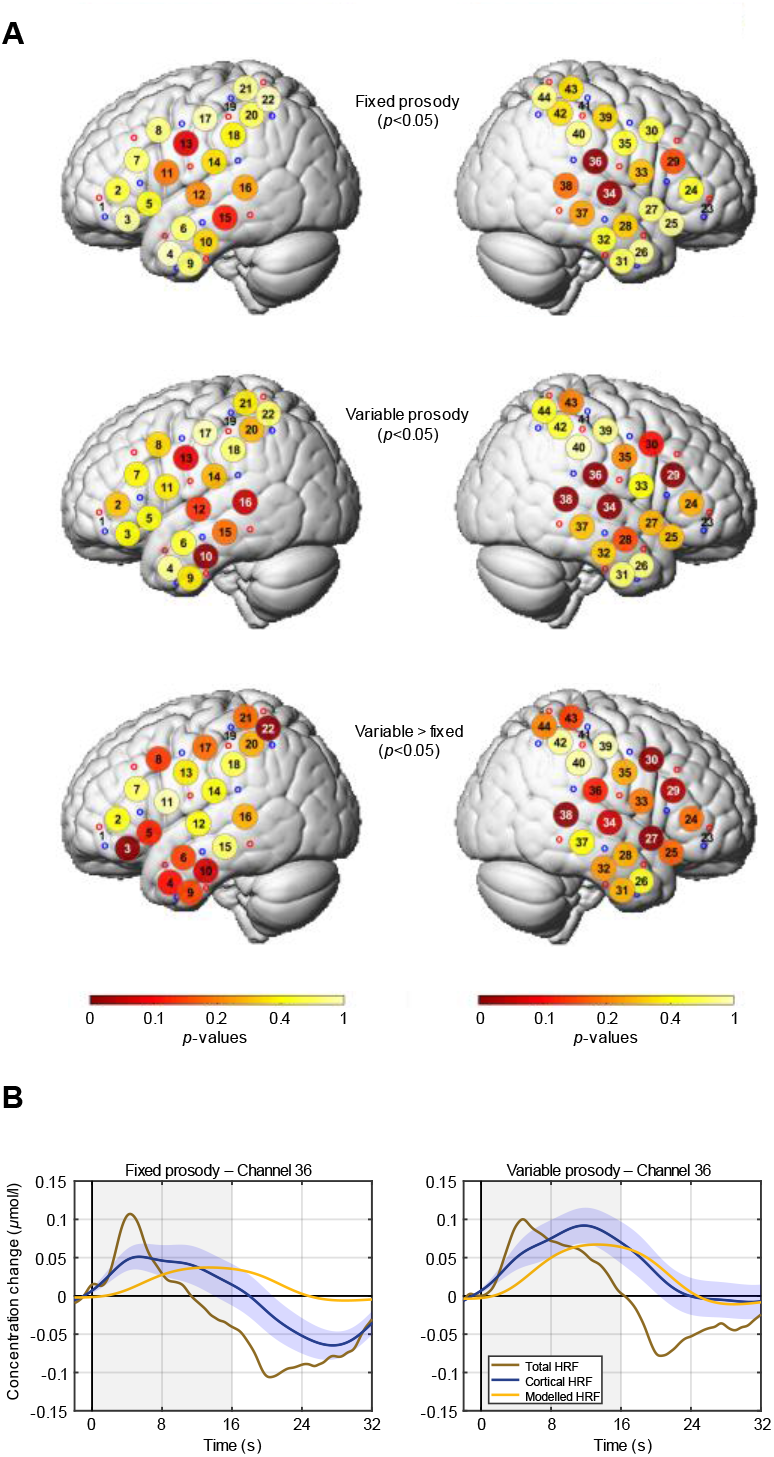
fNIRS oxygenated haemoglobin (HbO) results. A) Group-level fNIRS HbO topographies. The upper and middle rows show the activation maps for the conditions with fixed and variable prosody, respectively. The lower row shows the statistical comparison of both conditions. White channel numbers in the topographies indicate significance. B) Group-level HbO haemodynamic response functions (HRFs) for channel 36, located near the right auditory cortex, for the conditions with fixed and variable prosody. The shading around the cortical HRFs indicates the standard error of the mean.

### E. EEG recording and analysis

Continuous EEG signals were recorded using a BrainVision actiCHamp system (Brain Products, Gilching, Germany) with 60 electrodes arranged according to the extended international 10-20 system. Four additional electrodes were placed around the eyes to record vertical and horizontal eye movements. The EEG data were recorded with an initial sampling rate of 500 Hz, an online anti-aliasing low-pass filter with a cut-off frequency of 140 Hz and were referenced to the right mastoid. As with the fNIRS optodes, the electrode positions of each subject were digitized with a Polhemus 3SPACE ISOTRAK II system before the experiment.

The data were pre-processed offline using FieldTrip (version 20180924; Oostenveld et al., 2011) and custom MATLAB code. The continuous waveforms were first segmented into epochs ranging from −0.3–0.9 s relative to stimulus onset. Next, linear trends and the DC component were removed by subtracting a 1^st^-order polynomial and the epochs were low-pass filtered at 15 Hz (4^th^-order Butterworth, applied forwards and backwards). The epochs were then baseline corrected by subtracting the mean amplitude from −0.1–0 s before stimulus onset, re-referenced to the mean of both mastoids, and subsequently down-sampled to 250 Hz. After visually identifying and excluding bad channels (total = 9, max. 3 per subject), the data were decomposed into 20 principal components to detect and eliminate eye artefacts. After the 4 eye electrodes were removed from the data, epochs in which the amplitudes between −0.2–0.8 s around stimulus onset exceeded ±80 μV or the *z*-transformed amplitudes differed by more than 17.5 standard deviations from the mean of all channels were excluded from further processing. On average, 88% of the trials (1765/2000 per subject) passed the rejection procedure. In two subjects, EEG battery issues led to data loss, so that only 60% (1204) and 24% (479) of the trials remained after the rejection procedure. Lastly, bad channels were interpolated using the weighted average of the neighbouring channels and the data were re-referenced to the average of all 60 channels.

Grand-average event-related potentials (ERPs) were computed by averaging the epochs of each subject, applying a baseline correction from −0.1–0 s before stimulus onset, and calculating the group mean of the subject-level ERPs. The ERPs were statistically examined using non-parametric permutation tests (Maris and Oostenveld, 2007). To test if the ERP amplitudes were larger in the variable prosody condition compared to the fixed prosody condition, dependent-samples *t*-tests were calculated for all combinations of electrodes and sample points throughout the duration of the vowels (0–0.8 s). The scalp distributions of the ERP amplitudes, in contrast, were tested by first averaging the voltages over specific time windows. Here, false positive findings were controlled for by merging significant electrodes into spatially coherent clusters, if at least three neighbouring channels also showed the respective effect. The *t*-values in each cluster were then summed to derive a test statistic, which was compared to the same statistic after the trials were randomly re-allocated to the two conditions. This step was repeated 5,000 times to obtain the *p*-value of a cluster. For the ERPs, uncorrected tests were computed in the same way, but the individual *t*-values were re-allocated between groups to derive their *p*-values.

Distributed source reconstructions of the ERPs were computed using the MNE-dSPM approach implemented in Brainstorm (version 29-Apr-2020; Dale et al., 2000; Tadel et al., 2011). The electrode positions of each subject were first co-registered to the ICBM152 MRI template by aligning three external fiducial points (LPA, RPA, and Nz) and subsequently projecting the electrodes to the scalp of the template MRI. A Boundary Element Method (BEM) volume conduction model based on the ICBM152 template and the corresponding cortical surface (down-sampled to 15,000 vertices) were used as head and source models. The BEM head model was computed using OpenMEEG (version 2.4.1; Gramfort et al., 2010) and comprised three layers (scalp, outer skull, and inner skull) with 1082, 642, and 642 vertices, respectively. To validate the accuracy of the template-based source reconstructions, structural MRIs of four subjects were also obtained and used to devise individual head and source models. Linear MNE-dSPM solutions with dipole orientations constrained to be normal to the cortex were estimated after pre-whitening the forward model with the averaged noise covariance matrix calculated from the individual trials in a time window from −0.2–0 s before vowel onset. The default parameter settings for the depth weighting (order = 0.5, max. amount = 10), noise covariance regularisation (regularise noise covariance = 0.1), and regularisation parameter (SNR = 3) were used throughout. The individual source reconstructions were then converted to absolute values and spatially smoothed (Gaussian kernel, full width at half maximum = 3 mm). Statistical differences between the two conditions were evaluated using cluster-based non-parametric permutation tests, as described above.

## III. RESULTS

### A. fNIRS results

The grand-averaged fNIRS HbO topographies are shown in Fig. 3A. The upper two rows depict the functional activations for the fixed and variable prosody conditions compared to baseline level, whereas the lower row shows which channels showed stronger activity in the variable prosody condition compared to the fixed prosody condition. In both conditions, significant activity (*p*<0.05; indicated by white channel numbers) was observed around the right auditory cortex. However, as shown by the condition comparison, activity along the right superior temporal gyrus (channels 27, 34 & 38) was significantly stronger (*p*<0.05) in the variable prosody condition. Additional significant differences in the right hemisphere were observed in the pre-motor region (chs. 29 & 30). In the left hemisphere, in contrast, auditory cortex activity in both conditions was much less pronounced. No significant activity was observed in the fixed prosody condition, while two channels covering the temporal lobe (chs. 10 & 16) showed significant activity (*p*<0.05) in the variable prosody condition. Moreover, no significant differences between the two conditions were observed in auditory areas in the left hemisphere, while significance level (*p*<0.05) was reached for two channels in the inferior frontal and parietal cortex (chs. 3 & 22).

To complement the topographical results, the grand-averaged time course of the HbO concentration changes in the right auditory cortex are shown in Fig. 3B. For channel 36, covering the posterior part of the right auditory cortex, the cortical HRF in the variable prosody condition exhibited a markedly longer and stronger positive deflection compared to the cortical HRF observed in the fixed prosody condition. In particular, the cortical HRF in the variable prosody condition reached its peak towards the end of the stimulation period, as predicted by the canonical HRF model, while the cortical HRF in the fixed prosody condition already reached its peak during the beginning of the stimulation blocks and receded afterwards.

A qualitatively similar pattern of results was obtained for the analyses of the HbR data (Fig. 4). Here, the topographies of the individual conditions indicated somewhat more pronounced auditory cortex activity in both hemispheres compared to the HbO data, as reflected by a greater number of significant channels (*p*<0.05). The comparison of both conditions however only returned a trend for more activity along the right superior temporal gyrus in the variable prosody condition. Despite this lack of significance, the cortical HRFs of channel 36 closely resembled their HbO counterparts, with the variable prosody condition again showing stronger and more sustained concentration changes. Additionally, a small offset response after the end of the stimulus blocks is evident in the HRFs of both conditions.

**Figure 4.**
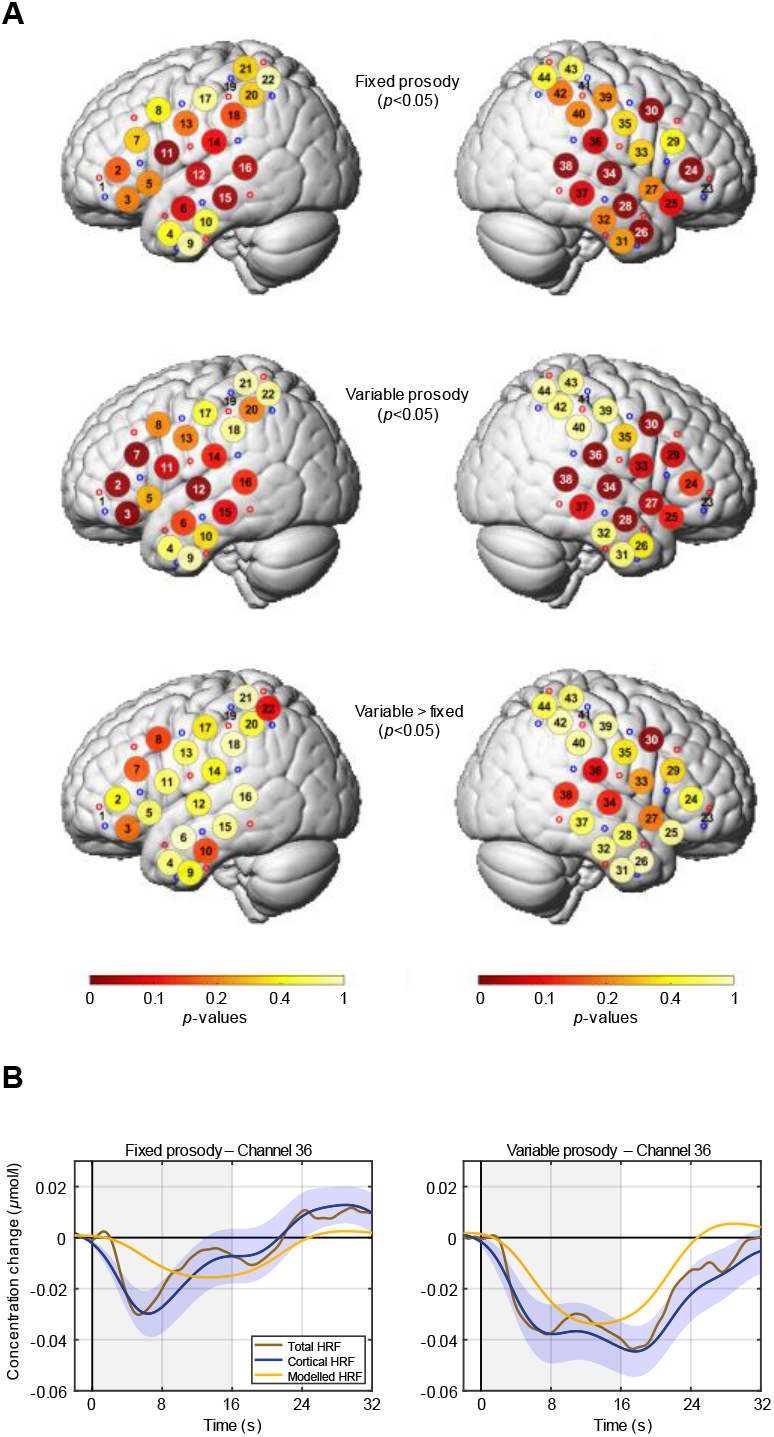
fNIRS de-oxygenated haemoglobin (HbR) results. A) Group-level fNIRS HbR topographies. B) Group-level HbR HRFs for channel 36. All plotting details are the same as in Fig. 3.

### B. EEG results

The grand-average ERPs as well as their statistical evaluation are shown in Fig. 5A. On the left, the sensor waveforms of electrode Fz are depicted for both conditions, together with their difference waveform (variable - fixed prosody). The statistical comparison of each time point throughout the duration of the vowels (0–800 ms) indicated significantly larger P1 and P2 amplitudes (*p*<0.001) in the variable prosody condition. The corresponding scalp maps are shown on the right of Fig. 5A. For both components, the time-averaged amplitudes in the variable prosody condition were found to be significantly larger (*p*<0.05, cluster-corrected) for a group of electrodes in the fronto-central scalp region.

**Figure 5.**
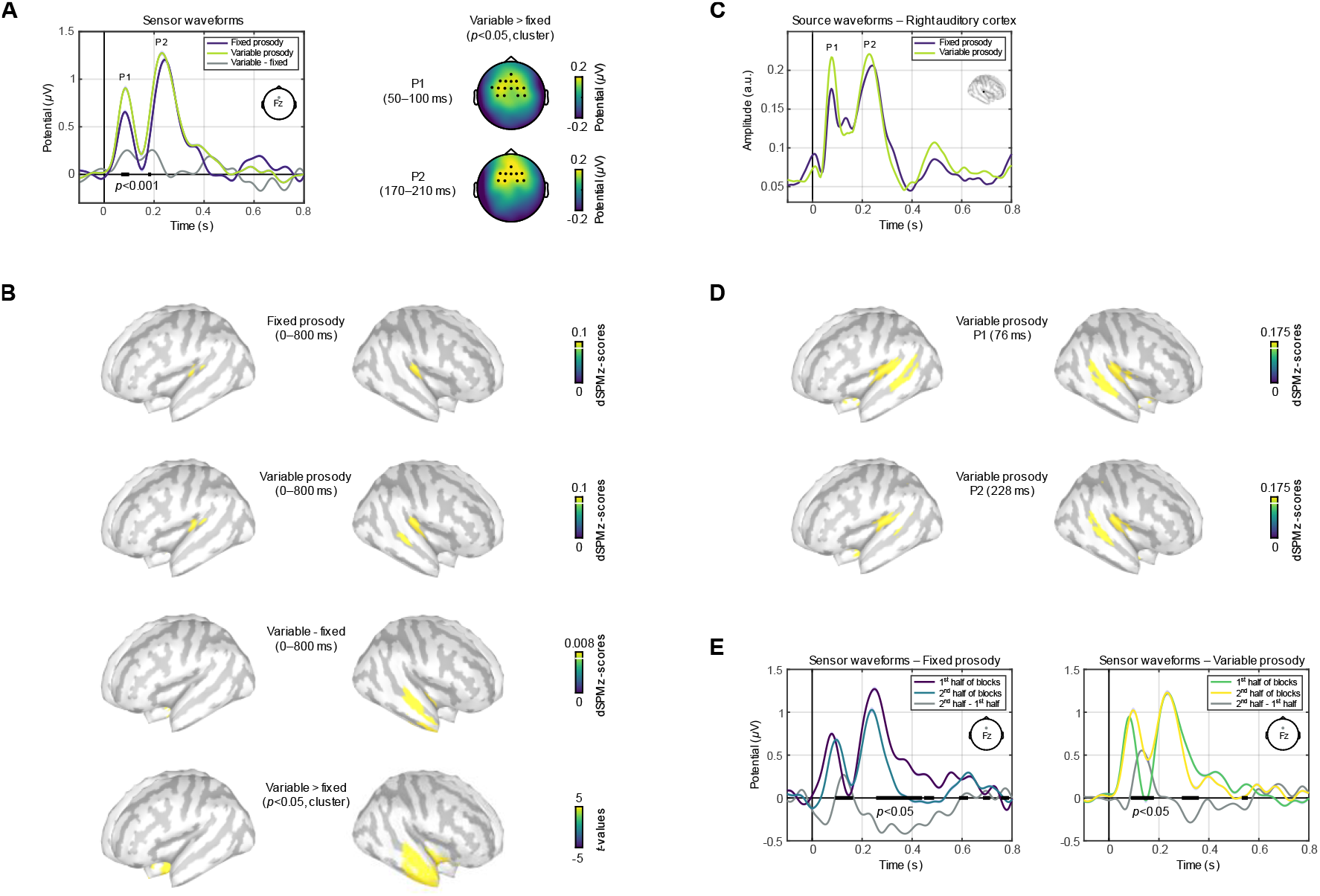
EEG results. A) Sensor-level event-related potentials (ERPs) and scalp maps. Significant differences in activity are indicated by the horizontal black bars below the ERPs. The shading around the ERP waveforms depicts the standard error of the mean. The scalp maps on the right show differences between conditions after time-averaging the voltages across the specified time windows. Electrodes for which the difference was significant are indicated by black dots. B) dSPM EEG source reconstructions averaged across the duration of the individual vowels. The upper two rows show the cortical activity for the fixed and variable prosody conditions, respectively. The third row exhibits the difference between the two conditions. Only cortical surface vertices with at least 85% of the maximal amplitude are plotted in colour. The lower row shows the statistical comparison of the two conditions. C) Group-level source waveforms extracted from the right auditory cortex (planum temporale). The surface location from which the waveforms were extracted is indicated by the black dot in the inlay plot. D) dSPM EEG source reconstructions for the P1 and P2 peaks in the group-level source waveforms for the variable prosody condition. E) Sensor-level ERPs for the first and second halves of the stimulus blocks in both conditions.

dSPM-based source reconstructions of the ERPs averaged across the whole duration of the vowels (0–800 ms) are shown in Fig. 5B. In both conditions, bilateral activity in the auditory cortex was observed, albeit strongly lateralised to the right hemisphere. In the variable prosody condition, there was additional activity in the right STS and activity in the right auditory cortex extended slightly further anterior. Subtracting the activity in the fixed prosody condition from that in the variable prosody condition revealed greater activity along the length of right STS as well as in the right planum polare in the variable prosody condition. These observations were confirmed by the statistical comparison of the two conditions, which returned one large cluster of vertices comprising right anterior temporal areas as well as the region anterior to the right auditory cortex that showed significantly larger activity (*p*<0.05, cluster-corrected) in the variable prosody condition.

To evaluate the accuracy of the source reconstructions, dSPM solutions of four subjects for which structural MRI scans were available were compared to solutions based on the ICBM152 template MRI. As shown in Suppl. Fig. 1, activity was generally larger when the subject-specific MRIs were used, while the overall right-lateralised pattern of activity remained unchanged. The individual MRIs thus enabled to localise a greater amount of the energy of the surface-recorded signals to the auditory cortex and adjacent regions but did not lead to results that were qualitatively different from the template-based solutions.

Group-level source waveforms reflecting the average time course of evoked activity at the source level are shown in Fig. 5C. These waveforms were extracted from a vertex point in the right planum temporale (MNI *x,y,z*-coordinates in mm: 41, −25, 16), the location that exhibited the strongest activity in both conditions. The source waveforms corroborate the ERP results shown in Fig 5A, as the P1 and P2 amplitudes were again larger in the variable prosody condition. In addition to the sensor waveforms, they also show increased sustained activity in the variable prosody condition during the second half of the vowels. Moreover, the source waveforms also exhibit a marked sound offset response, which was slightly larger in the fixed prosody condition.

In Fig. 5D, dSPM source reconstructions for the P1 and P2 peaks of the group-level source waveforms in the variable prosody condition are shown. The sources of the P1 in response to prosodic variations had a largely symmetric distribution, including the auditory cortex, planum polare, and STS in both hemispheres. For the P2, in contrast, bilateral activity was evident in the auditory cortex, whereas activity in STS and planum polare was largely confined to the right hemisphere.

Finally, based on the fNIRS results that indicated reduced response amplitudes in the fixed prosody condition towards the end of the stimulation blocks, we tested whether the ERP amplitudes differed between the first and second halves of the blocks. Due to its large energy-onset response, the first stimulus in each block was omitted in this analysis. As shown in Fig. 5E, the P1 amplitude did not adapt over the course of the blocks in both conditions, while the N1 was reduced in amplitude during the second half of the blocks in both conditions (*p*<0.05, two-tailed). The P2, however, showed a differential effect, with a more pronounced amplitude reduction during the latter half of the blocks in the fixed prosody condition compared to the variable prosody condition (*p*<0.05, two-tailed).

## IV. DISCUSSION

### A. Pitch changes elicit additional activity in right STS and planum polare

The present fNIRS and EEG results both indicated strongly right-lateralised activity in response to continuous vowel sequences in which the pitch contours of the individual vowels were either the same throughout the blocks (fixed prosody) or varied between the vowels (variable prosody). Crucially, both imaging methods also revealed additional activity along the right superior temporal lobe for sequences with variable prosody. The EEG source reconstructions furthermore revealed a smaller second region in the right planum polare, located anterior to the right primary auditory cortex, that showed stronger activity in the variable prosody condition. The present results are in line with previous studies that have found right-lateralised responses to prosodic variations, pitch changes, and melodies using fMRI (Meyer et al., 2002; Norman-Haignere et al., 2019; Patterson et al., 2002) and EEG source localisations of the activity evoked by consonant-vowel sequences (Dimitrijevic et al., 2013). In particular, Patterson et al. (2002) found increased right-lateralised activity to pitch changes in STS as well as in a second region slightly anterior to primary auditory cortex, in close agreement with the present EEG source reconstructions. The somewhat lower spatial resolution of the fNIRS data precluded such a distinction and the results instead indicated a single connected region along the superior temporal lobe that showed increased activity in the variable prosody condition.

### B. Strong right-lateralisation of the fNIRS and EEG results

Although the right-lateralisation in the present results was found to be particularly strong, it is consistent with the theoretical perspective that spectral variations are primarily processed in anterior superior temporal areas in the right hemisphere (Johnsrude et al., 2000; Zatorre et al., 2002) as well as the asymmetric sampling hypothesis (Boemio et al., 2005; Poeppel, 2003). The latter perspective explicitly states that the slowly varying prosodic features of speech sounds should lead to strongly right-lateralised processing. Furthermore, both frameworks assume that responses in primary auditory cortex should largely be symmetric and that only secondary areas in the superior temporal cortex should exhibit a strong lateralisation to the right when slow spectral changes in the stimuli are processed. Accordingly, this pattern was evident for the P2 in the variable prosody condition but not the preceding P1 (cf. Fig. 5D), demonstrating that this effect was not present during the initial stages of processing.

A methodological explanation for the overtly strong lateralisation may be that the 16-s stimulus blocks in the current experiment were longer than in previous studies and that the individual vowels within each block were not separated by silent gaps. Both of these factors serve to reduce energy-onset responses originating from the posterior part of primary auditory cortex in both hemispheres (Gutschalk et al., 2004; Krumbholz et al., 2003). Specifically, the studies by Dimitrijevic et al. (2013), Norman-Haignere et al. (2019), and Patterson et al. (2002) used stimulus sequences with durations ranging from less than 1 to 9 s and in two of these studies (Norman-Haignere et al., 2019; Patterson et al., 2002) the individual stimuli were separated by silent gaps.

### C. fNIRS and EEG responses to voice pitch changes are temporally sustained

The present fNIRS results showed that the HbO and HbR waveforms recorded near the right auditory cortex both exhibited not only higher amplitudes in the variable prosody condition, but also temporally more sustained responses. While the time course of these sustained responses matched the shape of the canonical HRF model well, the waveforms in the fixed prosody condition receded back to baseline level markedly earlier. This is in line with fMRI results showing that the haemodynamic response decreases after an initial peak when continuous, long stimulus blocks with a fixed pitch are presented (Steinmann and Gutschalk, 2012). However, while the higher temporal resolution of fNIRS generally allows for a more precise measurement of haemodynamic time courses, it has not been demonstrated yet that pitch changes serve to prolong the duration of the BOLD response.

Comparing the ERPs in the first and second halves of the stimulation blocks revealed that the sustained haemodynamic responses in the variable prosody condition appeared to be driven by a smaller adaptation of the P2 component. The P1, in contrast, exhibited similar amplitudes across both halves of the blocks in both conditions, whereas the N1 amplitudes were reduced in both stimulus conditions during the latter half of the blocks. The finding that only the P2 shows a differential adaptation effect in response to stimuli with fixed and variable pitch is also evident in previous data (Patterson et al., 2016; Rupp and Uppenkamp, 2005), but has not been explicitly investigated to date.

### D. Differences between the oxygenated (HbO) and de-oxygenated (HbR) fNIRS data

In accordance with the literature on the physiological bases of fNIRS signals (Kirilina et al., 2012; Tachtsidis and Scholkmann, 2016), the HbR data were less influenced by non-cortical artifacts, as indicated by the markedly greater similarity of the total and cortical HRFs. However, the smaller response amplitudes compared to the HbO data outweighed this advantage, resulting in topographies that were spatially coherent with the HbO results but showed smaller effects when contrasting the two conditions. Furthermore, the HbR time courses differed from their HbO equivalents as they contained separate sound onset and offset peaks, in line with fMRI data recorded in response to continuous auditory stimuli (Harms and Melcher, 2002; Steinmann and Gutschalk, 2012) and the causal relation of HbR and BOLD signal changes (Cui et al., 2011; Strangman et al., 2002).

In contrast to a previous fNIRS-EEG study in which we tested normal-hearing listeners with a comparable passive auditory paradigm (Steinmetzger et al., 2020), the HbO and HbR results were largely coherent in the current experiment. Specifically, our previous data showed prominent negative HbO responses in fronto-temporal areas, whereas the HbR data showed the expected activity in auditory areas. A possible explanation for this unexpected result might be that HbO signals are affected by activity of the sympathetic branch of the autonomic nervous system (ANS; Tachtsidis and Scholkmann, 2016). The absence of a similar effect in the current data thus suggests that no substantial changes in ANS activity were elicited.

Overall, the HbR data showed greater activity than the HbO data for the individual conditions, whereas the condition comparison instead returned greater differences for the HbO data. Beyond auditory areas, the fNIRS HbO and HbR topographies indicated stronger activity in the right motor cortex and the left inferior frontal gyrus (IFG) in the variable prosody condition. While passively listening to meaningless speech sounds has been shown to elicit activity in the motor cortex (Wilson et al., 2004), emulating the production of these sounds, this activity is often lateralised to the left (Pulvermüller et al., 2006). Silent singing, in contrast, has been found to predominantly activate the right motor cortex (Riecker et al., 2000). The stronger activity in the right motor cortex might hence be due to the song-like quality of the continuous pitch contours in the variable prosody condition. Moreover, activity in Broca’s area, the posterior part of left IFG, was shown to coincide with motor activity during speech perception (Watkins and Paus, 2004).

### E. ERP components associated with the processing of voice pitch changes

The variable prosody condition was found to elicit larger P1 amplitudes than the fixed prosody condition, a difference that was observed throughout the stimulus blocks. Despite a similar trend for larger P1 amplitudes in response to discrete pitch changes in melodies, compared to tone sequences with a fixed pitch, previous MEG studies (Patterson et al., 2016; Rupp and Uppenkamp, 2005) did not explicitly focus on this effect. However, the overall P1 amplitudes as well as the difference between conditions were also markedly larger in response to the vowel sequences in the current experiment.

The N1 amplitudes, in turn, were not found to differ significantly between the two conditions, in contrast to the results of Patterson et al. (2016) and Rupp and Uppenkamp (2005), where the N1 was larger and peaked slightly earlier in response to melodic pitch changes. As the fixed pitch condition in those studies consisted of the same tones in presented succession, the observed difference may be due to stronger adaption of the N1 amplitudes in that condition. Yet, in the fixed prosody condition in this experiment, the pitch changed continuously within the individual vowels, except for the flat contours. This likely resulted in less adaptation of the N1 and hence also a smaller relative difference compared to the variable prosody condition. Although the prosodic variations in the vowels sequences did not substantially affect the N1, the long stimulus blocks used in this study resulted in reduced overall N1 amplitudes compared to the earlier studies. The N1 was smaller during the second half of the vowel sequences and this adaptation effect was evident in both stimulus conditions.

The P2, on the other hand, was larger in the variable prosody condition, but this effect only emerged during the second half of the blocks. Similar P2 amplitudes across conditions were observed during the first half of the stimulus sequences, whereas the P2 was more strongly attenuated during the second half of the blocks in the fixed prosody condition. As the transient components of electrophysiological recordings have been shown to give rise to the BOLD response (Logothetis et al., 2001), this effect also seems to be the reason for the diminished fNIRS responses in the fixed prosody condition during the latter half of the stimulus blocks. Similarly, the consistently larger P1 amplitudes appear to underlie the stronger overall fNIRS responses in the variable prosody condition.

At the sensor level, no right-lateralisation was apparent for both the P1 and the P2, while the source reconstructions and the fNIRS results clearly showed this pattern. Likewise, no sustained negativity was evident in the ERPs at the sensor level because slow components with wide scalp distributions are strongly attenuated when an average reference is used and instead require a mastoid reference (Steinmetzger and Rosen, 2017). At the source level, however, the average time course of activity in the right auditory cortex showed increased sustained activity commencing about 400 ms after stimulus onset in the variable prosody condition. Although such sustained potentials do not drive the BOLD response (Gutschalk et al., 2010; Logothetis et al., 2001), and hence will also not have affected the current fNIRS results substantially, this finding is nevertheless in line with previous data. In the results of Patterson et al. (2016), the sustained field increased over the course of the melodies, but not when the same tone was repeated.

## V. CONCLUSIONS

The present fNIRS-EEG study has demonstrated that continuous vowel sequences in which the prosodic contours varied between the individual vowels evoke additional cortical activity in right STS and planum polare, anterior to the primary auditory cortex, compared to sequences in which the prosodic contours of all vowels were the same. The time courses of the fNIRS signals furthermore revealed sustained activity in response to sequences with changing prosodic contours, while sequences with fixed prosodic contours exhibited decreased activity during the second half of the stimulus blocks.

In the ERP data, a corresponding effect was evident for the P2 component evoked in response to the individual vowels, for which the amplitude in the condition with fixed prosody was markedly smaller during the latter half of the blocks. The P1 amplitude, on the other hand, was consistently larger for sequences containing prosodic changes and showed no such adaptation effect. The EEG source reconstructions of the P1 and P2 moreover indicated that the general right-lateralisation found in the fNIRS and EEG data was evident for the P2 but not the P1, and hence not present during the early processing stages.

More generally, the current results have shown that the spatial resolution of fNIRS data is sufficiently high to distinguish between the cortical representation of speech sounds with relatively subtle acoustic differences, at least in normal-hearing listeners. As the cortical activation maps obtained with fNIRS and EEG were largely coherent, the present data also serve as a mutual validation of both imaging methods. These results hence suggest that fNIRS is a suitable method for precise neurophysiological investigations of auditory perception, in addition to its many practical advantages.

## ACKNOWLEDGEMENTS

We are grateful to the Dietmar Hopp Stiftung (Grant No. 2301 1239) for supporting our research. Parts of this research have been presented at the *23^rd^ International Congress on Acoustics*: http://pub.dega-akustik.de/ICA2019/data/articles/000704.pdf.

## SUPPLEMENTARY MATERIALS

**Supplementary figure 1.**
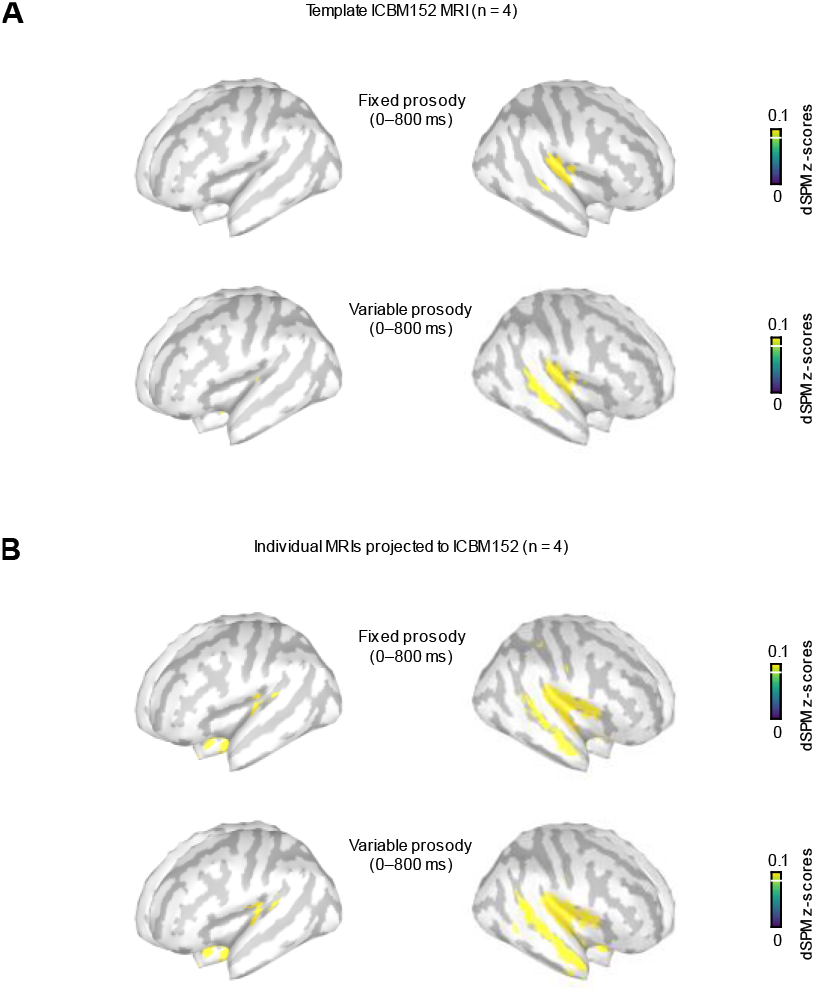
Averaged dSPM EEG source reconstructions of four subjects A) based on the template ICBM152 MRI and B) based on their individual MRIs and projected to the cortical surface of the ICBM152 template. All details, including the scaling, are the same as in Fig. 5B.

